# The contribution of the locus coeruleus – norepinephrine system to the coupling between pupil-linked arousal and cortical state

**DOI:** 10.1101/2025.05.07.652771

**Authors:** Evan Weiss, Yuxiang Liu, Qi Wang

**Affiliations:** Department of Biomedical Engineering Columbia University, New York, NY 10027, USA

**Keywords:** pupil-linked arousal, Locus Coeruleus, norepinephrine, neural oscillations, adrenergic receptors, EEG

## Abstract

Understanding how pupil-linked arousal couples with cortical state is crucial for uncovering the neural mechanisms underlying brain state-dependent cognitive and sensory processing. Pupil size fluctuations reflect rapid changes of the pupil-linked arousal system, indexing brain states as well as the activity of neuromodulatory systems, including the locus coeruleus-norepinephrine (LC-NE) system. We investigated the relationship among phasic pupil dilation, cortical state, and neuromodulation by combining optogenetic LC stimulation with EEG recordings and pupillometry in awake mice. A comparison between EEG signals during spontaneous phasic pupil dilation and those during phasic pupil dilation evoked by LC stimulation revealed distinct EEG power spectrums, with LC activation driving strong modulation in the alpha and beta bands. Using machine learning techniques, we trained a convolutional neural network classifier to distinguish between types of pupil dilation based on the power dynamics of individual EEG frequency bands. The results confirmed that EEG features in the alpha, beta, and gamma bands differ markedly between spontaneous phasic arousal and LC stimulation-evoked arousal. Moreover, pharmacological manipulations to either block α or β adrenergic receptors or agonize α-2 adrenergic receptors were employed to explore how adrenergic receptors could influence the coupling between phasic pupil dilation and cortical state. With each manipulation uniquely modulating EEG power and pupil size, our results highlight the differentiated role of adrenergic receptors in the maintenance of coupling between pupil-linked arousal and cortical state. This study provides new insights into the complex relationship between pupil-linked arousal and cortical arousal state, underscoring the significant role of the LC-NE system in influencing these arousal states.

**Significance Statement:** This study reveals the role of the locus coeruleus in pupil-linked arousal coupling with cortical arousal state by uncovering the different relationships between LC stimulation-evoked versus spontaneous phasic pupil dilations and cortical EEG. By integrating machine learning and noradrenergic pharmacological manipulation, our findings highlight the distinct cortical state associated with spontaneous phasic arousal and LC stimulation-evoked arousal, as well as the crucial role of different subtypes of adrenergic receptors in mediating the coupling between pupil size and cortical state.

## Introduction

The brain operates in various arousal states in our daily lives, which are regulated by central arousal systems and characterized by several physiological signals (Gilbert and Sigman, 2007; Niell and Stryker, 2010; Lee and Dan, 2012; Bennett et al., 2014; Poulet and Crochet, 2018). Historically, oscillation patterns in electroencephalogram (EEG) signals have been used to characterize cortical arousal state (Buzsáki and Draguhn, 2004; Jacobs et al., 2020). The changes in the distribution of power across different frequency bands (i.e., delta, theta, alpha, beta, gamma, or coarsely, low vs. high frequencies) have shown to correlate with transitions between wakefulness and sleep, and therefore are widely believed to reflect changes in arousal state (Steriade, 1996). Although other physiological signals, such as heart rate or heart rate variability, have been shown to index arousal state (Mathias and Stanford, 2003; Azarbarzin et al., 2014; Mather and Thayer, 2018; Wang et al., 2018; Liu et al., 2021), recent work has provided strong experimental evidence that fluctuations in pupil size are also a powerful indicator of the activity of a central arousal system (i.e. pupil-linked arousal) (Nassar et al., 2012; Murphy et al., 2014; Reimer et al., 2014; Ebitz and Platt, 2015; McGinley et al., 2015; Vinck et al., 2015; Urai et al., 2017; de Gee et al., 2020). This non-invasive measure of central arousal offers new insight into the effect of brain state on sensory and cognitive processing (Hess and Polt, 1960; Reimer et al., 2014; Reimer et al., 2016; Oliva and Anikin, 2018), as changes in pupil size have shown to provide a quantitative metric about shifts in attention, engagement, and internal models, making it a valuable tool for investigating how these changes influence neural coding and information processing (Gilzenrat et al., 2010; van Kempen et al., 2019; Zhao et al., 2019; Joshi and Gold, 2020; Schriver et al., 2020; Lapborisuth et al., 2022; Strauch et al., 2022).

Several lines of evidence have suggested that multiple neuromodulatory systems regulate pupil-linked arousal. Work by our group and others established the causal link between locus coeruleus (LC) activation and pupil dilation in both rodents and non-human primates (Joshi et al., 2016; Liu et al., 2017). In these studies, a burst of microstimulation of the LC mimicking phasic firing of LC neurons, evoked phasic pupil dilations. It has also been demonstrated that the activation of the dorsal raphe nucleus altered pupil size (Cazettes et al., 2021). Additionally, Reimer et al. (2016) showed strong correlations between pupil size and the activity of both noradrenergic and cholinergic axons in the cortex, consistent with a recent result where extracellular acetylcholine level in the prefrontal cortex was shown to co-vary with pupil size (Liu et al., 2024). It is important to note that the activation of these neuromodulatory nuclei also shift the power distribution across different frequency bands (Weiss et al., 2023). For instance, Liu et al. (2017) have demonstrated that phasic microstimulation of the LC of rats desynchronized cortical EEG activity by shifting EEG power from low-frequency bands (1–10 Hz) to high-frequency bands (10–100 Hz). Optogenetic stimulation of the LC increased EEG power in higher frequencies during NREM sleep periods and resulted in a significant decrease in theta activity (4– 9 Hz) during REM sleep in mice (Carter et al., 2010). Moreover, studies employing pharmacological manipulation of adrenergic receptors (α-1, α-2, and β) have showcased broad cortical state changes and behavioral consequences (Roubicek, 1976; Pastel and Fernstrom, 1984; Riekkinen Jr et al., 1990; Sainsbury and Partlo, 1993; Villain et al., 2016; Fitzgerald, 2021). Similar to LC stimulation, electrical microstimulation of the nucleus basalis, the cholinergic nucleus of the basal forebrain, increased the ratio of power in high-frequency bands relative to low-frequency bands of cortical LFP (Goard and Dan, 2009). As studies have demonstrated that spontaneous fluctuations in pupil-linked arousal are correlated with changes in cortical activity (Breeden et al., 2017), these neuromodulatory systems likely play a critical role in facilitating the coupling between pupil-linked arousal and cortical state (**Figure 1a**). However, little is known about the extent to which these neuromodulators contribute to the coupling.

**Figure 1:**
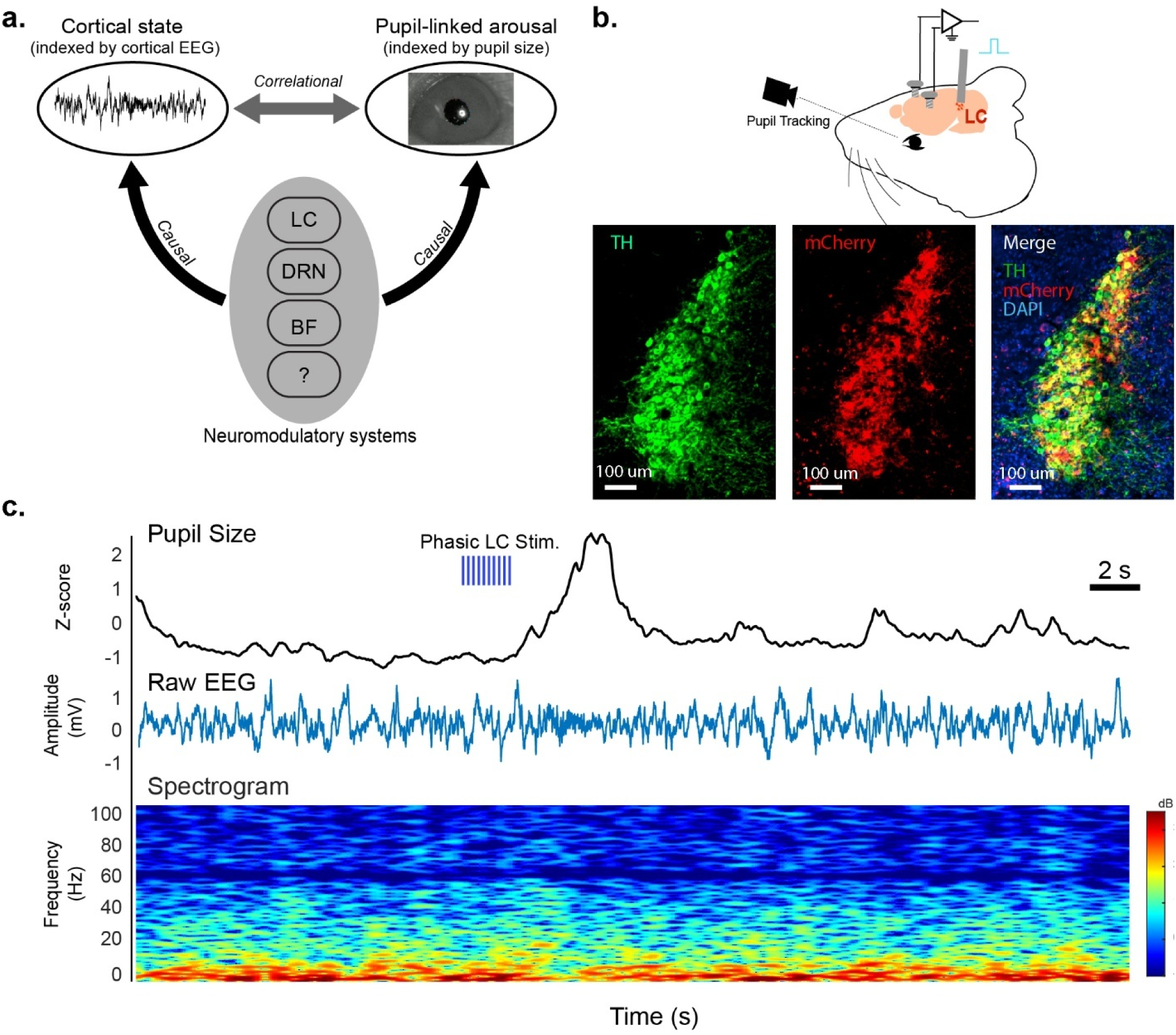
LC stimulation evokes pupil dilations and EEG spectral changes. **a.** Diagram of multiple neuromodulatory systems mediating coupling between pupil-linked arousal and cortical state. **b.** Diagram of simultaneous pupil and EEG recording during optogenetic LC stimulation (top), and histological confirmation of the expression of ChR2 in LC neurons (bottom). **C.** Visualization of a typical stimulation trial. Phasic pupil dilation evoked by phasic LC stimulation (top), raw EEG signal during stimulation period (middle), and an EEG spectrogram during same time period (bottom).

In this study, we aim to distinguish the role of the LC-NE system from the other neuromodulatory systems in mediating the coupling between phasic pupil-linked arousal and cortical state through optogenetic LC stimulation and pharmacological manipulation of different subtypes of adrenergic receptors. By quantifying the difference between cortical state during spontaneous phasic pupil dilation periods and LC stimulation-evoked phasic pupil dilation periods, we found distinct EEG power dynamics across the six canonical frequency bands during the two types of phasic pupil dilations. To further quantify differences, we developed a convolutional neural network classifier to decode the type of pupil dilation from EEG power across each individual frequency band, and confirmed that spontaneous and LC stimulation-evoked phasic pupil-linked arousal exert distinct effects on EEG power dynamics, particularly in higher frequency bands. Furthermore, we employed pharmacological manipulation of adrenergic receptors using propranolol, phentolamine, and clonidine to explore how the manipulation of each subtype of adrenergic receptors affected pupil-cortical state coupling. We found that each drug uniquely altered underlying spontaneous pupil dynamics as well as power distribution across EEG frequency bands. Optogenetic LC stimulation during each pharmacological manipulation seemingly did not compensate for the widespread effects across cortical EEG, indicating an important role of each subtype of adrenergic receptor in the maintenance of pupil-cortical coupling. Taken together, these results provide new insights into the complex coupling between pupil-linked arousal and cortical state, highlighting the unique influence of the LC and adrenergic receptors in the coupling.

## Materials and Methods

All experimental procedures involving animals were approved by the Columbia University Institutional Animal Care and Use Committee (IACUC) and were conducted in compliance with NIH guidelines. Adult Dbh-Cre mice (RRID: IMSR_JAX:033951, Jackson Lab) of both sexes (9 mice, 5 females), aged 3 ∼ 7 months, were used in the experiments. All mice were kept under a 12-hour light-dark cycle.

### Surgical Procedures

In aseptic setup, mice were anesthetized with isoflurane in oxygen (5% induction, 2% maintenance) and fixed in a stereotaxic frame. Body temperature was maintained at 36.6 ℃ using a feedback-controlled heating pad (FHC, Bowdoinham, ME). Following the removal of fur from the scalp and cleaning of the surgical site, lidocaine (0.1 ml, SC) was injected to the scalp to provide local anesthesia. Buprenorphine (0.05 mg/kg, sc) was then administered to ensure analgesics were on board throughout the surgery.

For adeno-associated viral vector (AAV) injections, a burr hole was drilled above the left LC. Pulled capillary glass micropipettes (Drummond Scientific, Broomall, PA) were back-filled with AAV solution and injected into the target brain regions at 0.7nL/s using a precision injection system (Nanoliter 2020, World Precision Instruments, Sarasota, FL). The pipette was left in place for at least 10 minutes between injections and slowly withdrawn. To optogenetically activate the LC, pAAV-EF1a-double-floxed-hChR2-(H134R)-mCherry-WPRE-HGHpA (Addgene #: 20297- AAV9, 250 nL) was injected into the LC (AP: -5.3 mm, ML: 0.85 mm, DV: -3 mm). Immunohistology staining for tyrosine hydroxylase (TH) confirmed the expression of mCherry-tagged ChR2 in TH-positive neurons in the LC (**Figure 1b**). After injection, an optical fiber (200μm diameter & NA = 0.39) was implanted with the tip of the fiber placed approximately 150 μm above injection site. C&B Metabond (Parkell Inc., Edgewood, NY) was used to build a headcap to bond the ferrule and the head bar.

For EEG electrode implantation, three burr holes were drilled above the prefrontal cortex (bilaterally, ML: ±0.4 mm, AP: 1.5 mm) and occipital lobe (ML: -0.5 mm, AP: -4.0 mm), with saline applied to each craniotomy to prevent drying out of brain surface. Three EEG recording screw electrodes (NeuroTek-IT Inc, Toronto, Canada) were then threaded into the skull to gently touch the brain surface. The wire of the EEG screw electrodes was connected to a custom-made EEG head stage with conductive epoxy (MG Chemicals 8331S-15G). The head stage was then bonded to the headcap with light-curing dental cement (Prime Dental Manufacturing, Chicago, IL). At the conclusion of the surgery, Baytril (5 mg/kg) and ketoprofen (5 mg/kg) were administered. Four additional doses of Baytril and two additional doses of ketoprofen were provided every 24 hours after the surgery day. All recordings were performed at least 3 weeks after surgery to allow sufficient time for viral expression.

### Pharmacological Manipulation

The pharmacological manipulation of the LC-NE system was done through administration of three FDA-approved adrenergic drugs: propranolol hydrochloride (Spectrum Laboratory Products PR 140), a non-selective beta blocker (Villain et al., 2016), phentolamine hydrochloride (Sigma-Aldrich P7547), an α-adrenoceptor antagonist (Starke et al., 1971; Cazala, 1980), and clonidine hydrochloride (Spectrum Laboratory Products CL118), an α-2 adrenergic agonist (Delbarre and Schmitt, 1971; Kleinlogel et al., 1975; Nelson et al., 1985). Each drug was dissolved in saline and sterile filtered before use and was administered intraperitoneally (IP) approximately 15-30 minutes before sessions. Dosage in mg/kg was calculated based on experimental animals’ daily weight. Propranolol was dosed at 1 mg/kg, Phentolamine at 5 mg/kg, and Clonidine at 1 mg/kg. Saline control sessions were randomly interleaved daily during experimental recording periods.

### Histology

At the end of the study, mice were transcardially perfused with PBS followed immediately by ice-cold 4% paraformaldehyde. The brain was removed carefully and post-fixed overnight at 4 °C in 4% paraformaldehyde, and then cryopreserved in a 30% sucrose (wt/vol) in PBS solution for 3 days at 4 ℃. Brains were embedded in Optimum Cutting Temperature Compound, and 25- μm coronal slices were sectioned using a cryostat. Brain slices were washed 4x in PBS and then incubated in 10% normal goat serum contained with 0.5% Triton X-100 in PBS for 2 hours. This was followed by primary antibody incubation overnight at room temperature using a chicken anti-TH (1:500) primary antibodies. On the next day, slices were washed 3x in PBS + Tween (0.0005%) solution followed by secondary antibody incubation for 2 hours at room temperature using an Alexa Fluor 488-conjugated goat anti-chicken (1:800). The slices were then washed 3x in PBS + Tween solution and 1x with PBS only followed by cover slipping using Fluoromount-G medium with DAPI. Selected example slices were imaged using 10X under a confocal microscope (Nikon Ti2) with a spinning disk (Yokogawa CSU-W1).

### Optogenetic Stimulation and EEG recording

To photo stimulate LC neurons expressing ChR2, blue light generated by a LED module was delivered through the implanted optical fiber (λ = 473 nm, PlexBright, Plexon Inc., Dallas, TX). During recordings, the mouse sat in a 3D printed head-fixation platform. Every 30 seconds, an optogenetic stimulation with either 5 or 10 Hz (randomly interleaved,10ms pulse duration) was delivered for a 2-second duration. EEG signals were digitalized by a 16-channel recording head stage (Model #C3335, Intan Technologies) and subsequently recorded by a RHD USB interface board (Model #C3100, Intan Technologies). A Bpod State Machine (Sanworks) was used with custom Python experimental scripts to synchronize optogenetic stimulation events, EEG recording, and pupil camera acquisition.

### Pupillometry and Pupil Size Extraction

Pupil recordings were obtained using a custom pupillometry system (Liu et al., 2024). The camera was triggered by 10 Hz TTLs from the Bpod State Machine. Pupil images were streamed to a high-speed solid-state drive for offline analysis. For each video clip, a region of interest (ROI) was manually selected initially. The DeepLabCut toolbox was used to segment the pupil contour (Mathis et al., 2018). Training sets were created, consisting of 90 frames for video clips with the resolution of 1280*1080 pixels. Within each frame, 12 points around the pupil were manually labeled, and cropping parameters were adjusted to enhance training accuracy. The resnet_50 deep network was trained on each frame and employed for the analysis of video clips from all sessions. Circular regression was then applied to fit the automatically labeled points, enabling the computation of pupil size based on the fitted contour. To ensure segmentation accuracy, approximately 5% of segmented images were randomly selected and inspected. Pupil size during periods of blinks was estimated through interpolation, using pupil sizes just before and after blinks. Prior to further analysis, a fourth-order non-causal low-pass filter with a cutoff frequency of 3.5 Hz was applied to the pupil size data (Schriver et al., 2018).

### Data Analysis

All data analyses were first conducted on individual sessions. Grand averages and standard errors of means were then calculated across sessions or animal subjects for analysis and visualization.

### Spontaneous Pupil Dilation Detection

Pupil size of each session was first z-scored and then subsequently up-sampled from 10 Hz to 100 Hz using spline interpolation. Up-sampling was necessary to match our EEG power time series to our pupil data for cross-correlation analyses. Next, a high-pass non-causal Butterworth filter was used to remove low-frequency drifts from the pupil data with a cutoff frequency of 0.1 Hz. To detect spontaneous pupil dilations, the following protocol and criteria were used for consistency across sessions and animals: the pupil size time series was iteratively segmented into 10-second windows, sliding through the entire time series for each session in 0.01 second increments. For each window, a baseline stability period was confirmed by checking whether the first 2 seconds remained continuously below a z-score threshold of 1, ensuring the pupil size was stable before detecting a peak. If baseline criterion was met, the algorithm searched for peaks within the same 10 s window where the z-score exceeded 1 (i.e., 1 standard deviation). The first detected peak was identified as the max of the dilation event. After the peak, the algorithm checked whether the pupil returned to baseline (below the baseline z-score threshold) within 6 seconds, ensuring short, transient dilations were captured. Each dilation was bounded by the period preceding the first peak and the interval following it. Spontaneous dilations events were aligned by their peak, providing a common reference point and a central point treated as time 0 across trials (**Figure 3a**). Once a dilation was identified, the algorithm will search for the next spontaneous dilation from the offset of the current dilation, and this process was been repeated through the recording of the entire session.

For LC stimulation-evoked dilations, the alignment windows were also aligned by the peak of the pupil dilation, identical to the spontaneous method described above. Comparing this method to an alignment of LC stimulation-evoked dilations to stimulation onset, yielded a similar pupil dilation in peak dilation and baseline size (**Supplemental Fig. 2a**). This approach ensured identically sized and aligned temporal windows, allowing for a fair comparison of the time course and magnitude of the pupil responses across spontaneous and LC stimulation-evoked conditions.

### EEG Processing and Analysis

The recorded EEG was first filtered with a non-causal 6th order low-pass Butterworth filter with a cutoff frequency of 100 Hz to remove high-frequency noise. To align the 100 Hz pupil time series with the 1000 Hz original sampling frequency of the EEG data, we used a sliding window technique. This down sampled the EEG data to match the time resolution of the pupil data, ensuring both signals were temporally aligned and suitable for analysis. The moving window analysis was implemented by segmenting the EEG data sampled at 1000 Hz into 1-second windows, each containing 1000 samples. To ensure smooth temporal coverage, the step size was set to 10 samples (10 ms), ensuring the resulting EEG power series had the same 100 Hz as the pupil data. For each sliding window, we computed its power spectral density (PSD) using the Fast Fourier Transform (FFT) of the EEG signal within that window (Delis et al., 2018). To evaluate the power within each frequency band, we computed the area under the curve (AUC) of the PSD for specific frequency bands: Delta (1-4 Hz), Theta (4-8 Hz), Alpha (8-12 Hz), Beta (12-30 Hz), Low Gamma (30-55 Hz), and High Gamma (65-100 Hz). The AUC for each frequency band was calculated using MATLAB’s *trapz* function, approximating the integral of the PSD curve, yielding average power within that frequency range. This process was repeated every 10 ms across the entire dilation period, producing a PSD time series that matched the resolution of the pupil data (**Figure 4a**).

### Convolutional Neural Network Classifier Analysis

To classify the types of phasic pupil dilation from EEG band power, a convolutional neural network (CNN) was applied. For each frequency band, the PSD time series data, temporally centered around peak pupil dilation, was processed. This included the construction of a dataset which included all trials from all experimental sessions across all animals. A separate neural network model, with the architecture detailed below, was individually trained and evaluated for each frequency band. Model performance was assessed using a 5-fold-cross-validation scheme. Each model was trained using the Adam optimizer with a minibatch size of 32 sequences. Training was conducted for a maximum of 100 epochs. An early stopping criterion, monitoring the validation accuracy was employed to prevent overfitting and ensure that the model weights from the epoch yielding the highest validation accuracy were retained. Our framework was implemented in Python using the TensorFlow library with the Keras API. All training and validation were executed on an NVIDIA A100 GPU.

Our CNN employed ReLU activation, adjusting a feature space for each layer:

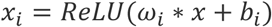

where *ω*_i_ are the filters, * denotes convolution operation, *b*_i_ are the biases, *x*_i_ is the output feature map after activation.

The network’s output followed a single unit dense layer with sigmoid activation function to output a probability indicating the class label, using the following activation:

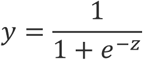

where z is the input into the sigmoid function and y is the output probability.

### Statistics

All statistical tests were two-sided. If the samples were normally distributed, a paired or unpaired t-test was used. For comparisons involving more than two groups, a one-way ANOVA was conducted, followed by a post hoc Tukey test to account for multiple comparisons. If the data did not meet the assumptions of normality, a Mann-Whitney U test was employed as a nonparametric alternative for comparisons between two groups. A Wilcoxon rank-sum test was used as a non-parametric alternative for comparison between two groups when normality was violated.

## Results

### LC stimulation evokes pupil dilations and EEG spectral changes

While previous behavioral studies have quantified the LC’s ability to modulate arousal, a clear investigation into its role in mediating the relationship between pupil-linked arousal and cortical state is missing. To address this question, we employed optogenetic stimulation to selectively activate LC neurons, while simultaneously recording pupil and EEG activity (**Fig. 1b**). In Dbh-Cre mice we injected a pAAV-EF1a-double-floxed-hChR2-(H134R)-mCherry-WPRE-HGHpA viral vector to drive Channelrhodopsin (ChR2) expression specifically in LC neurons, activating them by delivering blue light through an optical fiber implanted above the LC. While optically stimulating the LC, we simultaneously recorded the animal’s pupil size and EEG signals via the implanted skull electrodes (**Fig. 1b**). Post-mortem immunohistochemistry staining verified the ChR2 expression in the LC, showing robust overlap between mCherry, a tag of the AAV, and

TH signals, a hallmark of LC neurons (**Fig. 1b**). We also observed phasic pupil dilations and changes in EEG spectrum following optogenetic LC stimulation (**Fig. 1c**).

To further assess the distinct effects LC stimulation has on cortical EEG, we compared EEG spectrum between LC stimulation sessions and non-stimulation control sessions to quantify the extent to which elevated LC activity affects cortical arousal state. In each mouse we recorded multiple spontaneous (i.e., no LC stimulation) sessions as well as LC stimulation sessions, allowing us to compute the average normalized power spectral density across both session types (**Fig. 2a**). Consistent with previous results showing that LC-driven shifts in cortical arousal, here we observed a notable decrease in power located in low frequencies, with opposing increases in high frequency power in LC stimulation sessions compared to control sessions (Berridge and Foote, 1991; Bouret and Sara, 2005). Comparison between spectra of raw EEG signals in sessions with and without LC stimulation exhibited similar trend (**Supplementary Fig. 1**). Further quantifying EEG power band by band confirmed significant increases in low gamma (p=3.2792e- 05, Mann-Whitney U test, **Fig. 2b**) and high gamma power (p=5.6348e-10, Mann-Whitney U test, **Fig. 2b**), as well as a significant decrease in alpha power (p=7.6566e-04, Mann-Whitney U test, **Fig. 2b**).

**Figure 2:**
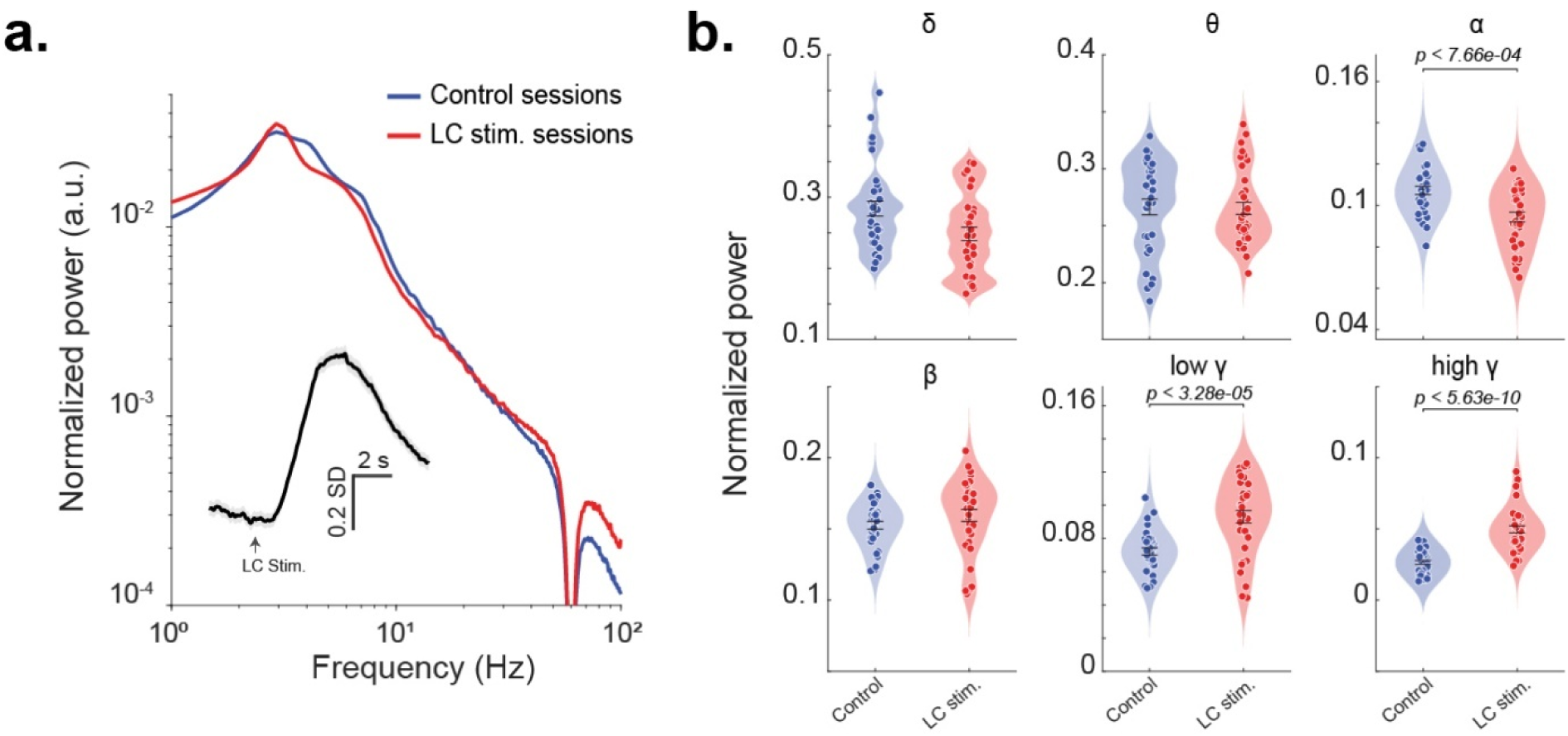
EEG spectrum of LC stimulation and control sessions. **a)** Normalized power spectral density spectrum between spontaneous and LC stimulation sessions, Inset: pupil size aligned to LC stimulation onset. **b)** Normalized power over each frequency band (δ (1–4 Hz), θ (4–8 Hz), α (8–12 Hz), β (12–30 Hz), low γ (30–55 Hz) and high γ (65–100 Hz)) for LC stimulation sessions and control sessions.

As previous work suggests that the activation of other neuromodulatory systems other than the LC-NE system can affect pupil size, these neuromodulatory systems may also contribute to the coupling between pupil-linked arousal and cortical state. To better understand the role of the LC-NE system in mediating the coupling between pupil-linked arousal and cortical state, we compared cortical EEG signals during spontaneous phasic pupil dilation periods and LC-stimulation evoked pupil dilation periods. To this end, we first identified spontaneous phasic pupil dilations (see Methods). On average, spontaneous phasic dilations occurred every 36.06 s, which is comparable to intervals of LC stimulations (**Fig. 3a inset**). However, unlike LC stimulation evoked pupil dilations, the onset of spontaneous phasic pupil dilations is hard to measure. Therefore, we aligned these two types of dilations by their peaks (**Fig. 3a**). Notably, aligning LC stimulation-evoked pupil dilations by the onset of optogenetic stimulation did not significantly differ from these pupil dilations aligned by their peaks (**Supplemental Fig. 2**). Although spontaneous phasic pupil dilations had a smaller baseline than LC stimulation-evoked pupil dilations (p=0.0140, Student’s t-test, **Fig. 3b**), their amplitudes were significantly larger (p=2.6106e-04, Student’s t-test, **Fig. 3b**).

**Figure 3:**
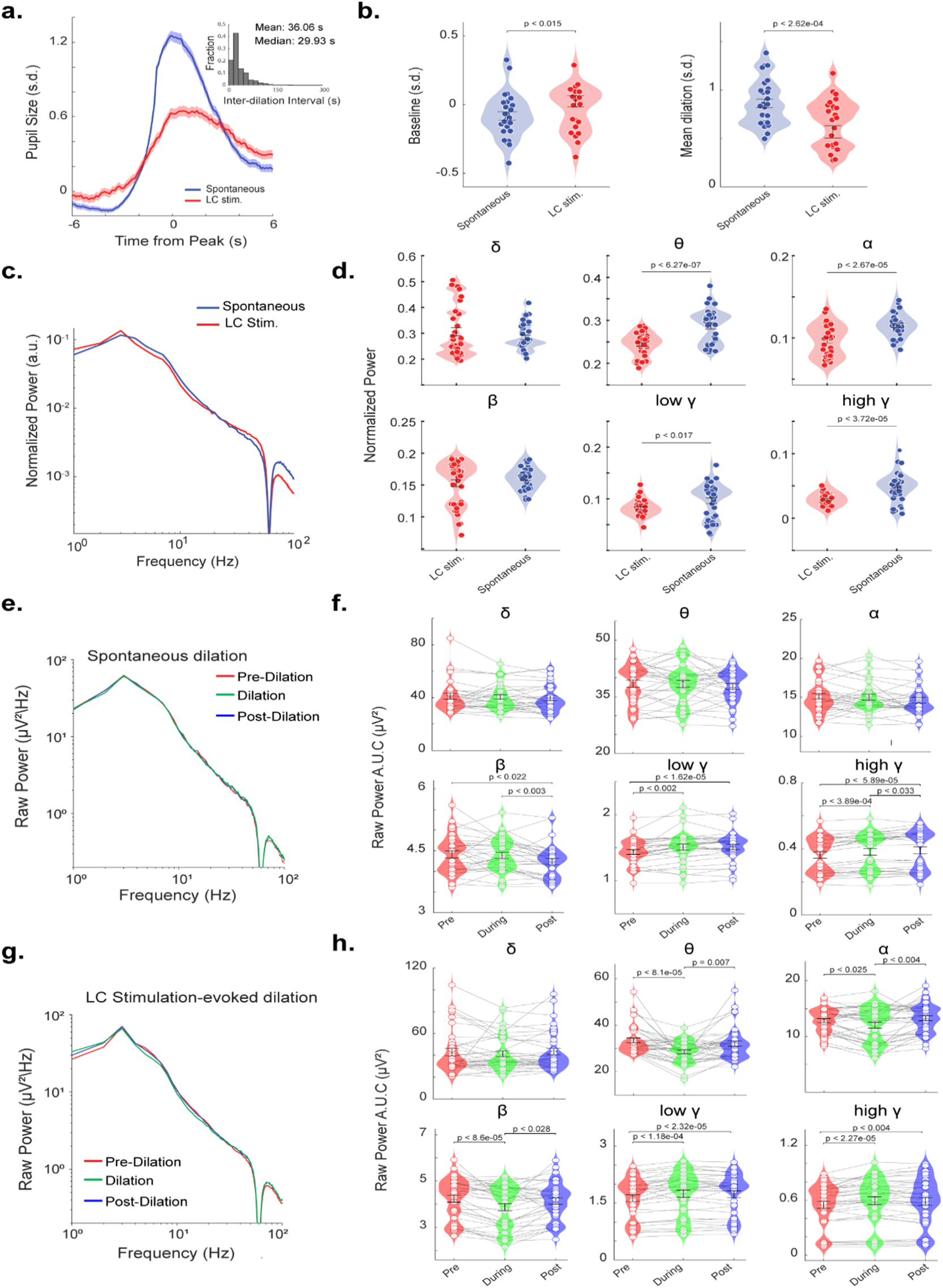
EEG spectrum around spontaneous and LC stimulation-evoked phasic pupil dilations. **a.** Pupil size aligned to peak dilation (-6 to 6 s from peak) for spontaneous and LC stimulation-evoked pupil dilations. Inset: histogram of inter-dilation intervals of spontaneous phasic dilations; mean time between dilations: 36.06 s, median: 29.93s. **b.** Quantification of pupil baseline (left) and mean dilation size (right) for spontaneous and LC stimulation-evoked phasic pupil dilations. **c, d.** Normalized EEG power spectrum and power in each frequency band during spontaneous and LC stimulation-evoked pupil dilations. **e, f.** Raw EEG power spectral density and power in each frequency band before, during and after spontaneous phasic pupil dilations. **g, h.** Raw EEG power spectral density and power in each frequency band before, during and after LC-stimulation evoked phasic pupil dilations.

We next looked to quantify different EEG power distributions during LC stimulation-evoked versus spontaneous pupil dilations. Computing the normalized EEG power spectrum during each type of pupil dilation periods revealed a distinct difference in low and high frequency band (**Fig. 3c**). Notably, EEG signals during LC stimulation-evoked pupil dilation periods had significantly lower power in the theta (p=6.2698e-07, Student’s t-test, **Fig. 3d**), alpha (p=2.6601e-05, Student’s t-test, **Fig. 3d)**, low gamma (p=0.0168, Student’s t-test, **Fig. 3d**), and high gamma (p=3.719e-05, Student’s t-test, **Fig. 3d**) bands.

After identifying different cortical states associated with spontaneous pupil dilations and LC stimulation evoked pupil dilations, we next compared cortical states right before, during, and after each of the two types of pupil dilations to gauge the time course of the coupling between pupil-linked arousal and cortical state. To this end, we calculated EEG spectrum within three 5- second periods, i.e. [-7.5 -2.5], [-2.5 2.5], and [2.5 7.5] seconds relative to the peak of pupil dilations. During spontaneous pupil dilations, there was a significant increase in power in both the low and high gamma bands compared to pre-dilation periods (p=0.0014 and p=3.8881e-04, respectively, paired t-test, **Fig. 3f**). On the contrary, in LC stimulation conditions, we observed significant changes in power in all bands except the delta band between the dilation periods and the pre-dilation periods. Specifically, there was a decrease in power in the theta, alpha, and alpha bands (p=8.0443e-05, p=0.025, and p=8.5629e-05, respectively, paired t-test, **Fig. 3h**), and increase in power both the low and high gamma bands (p=1.1725e-04, and p=2.2636e-05, respectively, paired t-test, **Fig. 3h**). Together, this difference in pupil-cortical state coupling between spontaneous pupil dilations and LC stimulation-evoked pupil dilations suggests the involvement of other neuromodulatory systems in mediating the coupling between pupil-linked arousal and cortical state.

### Convolutional neural network classifier analysis confirmed distinct power dynamics across the EEG bands during LC stimulation-evoked and spontaneous pupil dilations

Given spontaneous pupil dilations and LC stimulation-evoked pupil dilations are associated with different EEG power spectra, we looked to further quantify the temporal dynamics of pupil-cortical state coupling. To visualize EEG spectral power variations around phasic pupil dilation periods, we computed power spectral density using a moving window. This offered higher resolution into band-specific power dynamics over time, which were not measurable with the prior dilation windowing approach (**Fig. 4a**). In this method, EEG power during both LC stimulation-evoked and spontaneous pupil dilations exhibited clear drops in low-frequency (delta and theta) power as well as increases in high-frequency (low and high gamma) power. This confirmed the relationship described earlier, where both types of dilations were associated with the shift of cortical state from deactivated (higher power in low-frequencies) to activated (higher power in high-frequencies states). Interestingly, while power in the alpha band decreased significantly before LC stimulation-evoked pupil dilations reached their peaks, there was no obvious change in alpha band power during the same periods for spontaneous pupil dilations (**Figure 4a**).

**Figure 4:**
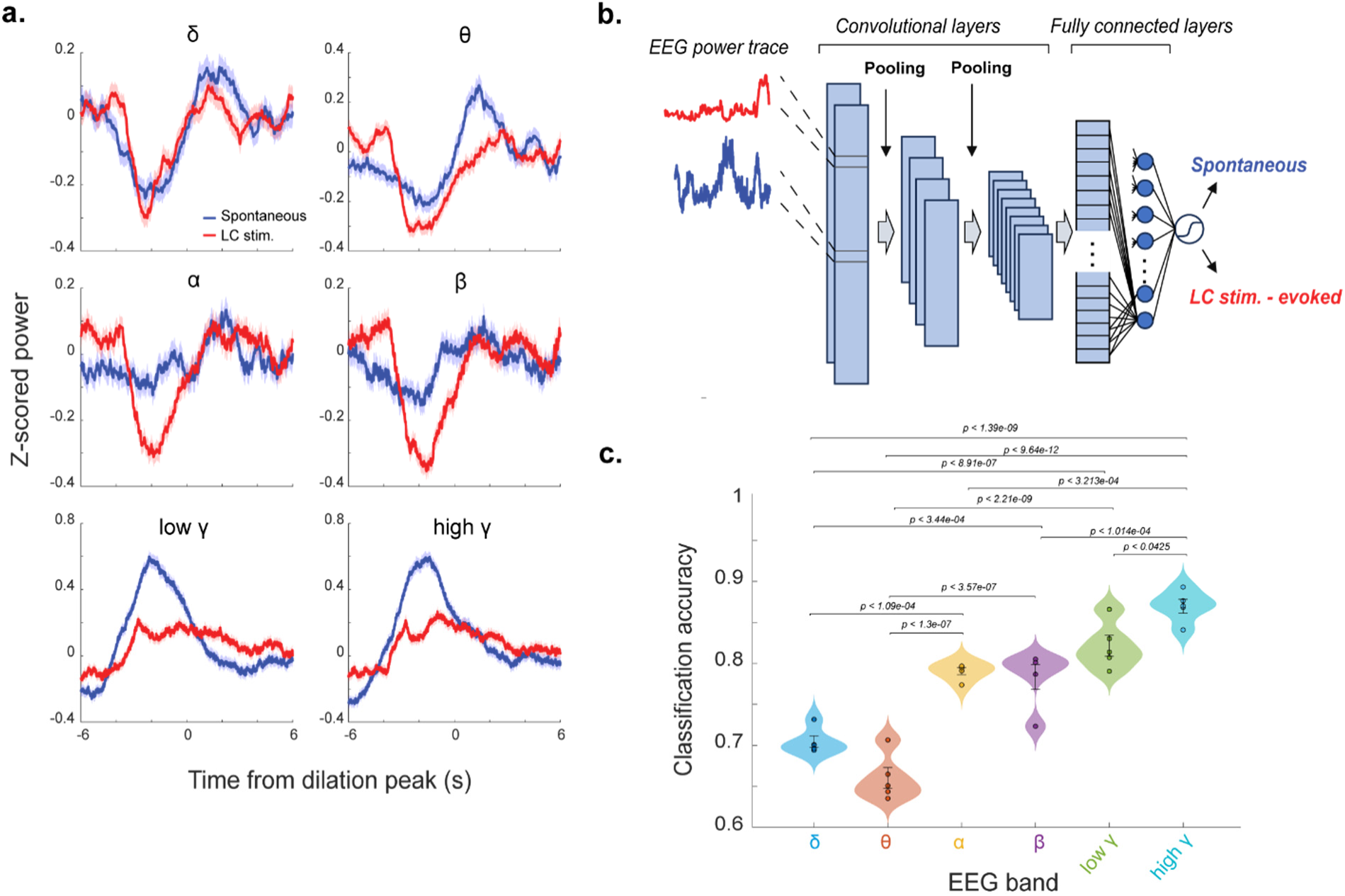
CNN classification of type of pupil dilations from EEG power dynamics. **a.** EEG power dynamics of each EEG frequency band around spontaneous and LC stimulation-evoked phasic pupil dilations. **b.** Architecture of the convolutional neural network. EEG power traces were fed into a convolutional neural network with multiple convolutional, fully connected, and pooling layers with sigmodal output being the network’s estimation of a specific dilation being spontaneously evoked or from LC stimulation. **c.** Model classification accuracy across training folds for each frequency band.

In order to further elucidate the relationship between pupil-linked arousal and cortical state under spontaneous and LC stimulation conditions, we looked to machine learning to help classify the hidden dynamics in each EEG spectral band during LC stimulation-evoked and spontaneous pupil dilations. We developed a convolutional neural network (CNN) classifier to decode if a pupil dilation is from spontaneous elevation of arousal or evoked by LC stimulation using each frequency band’s power spectral time series (**Fig. 4b**). In training, the network was fed frequency band-specific EEG power traces associated with the two types of pupil dilations across all animals, passing them through multiple convolutional layers to generate a single sigmoidal output. Our CNN decoder was evaluated using 5-fold cross-validation, giving us an average classification accuracy to predict the type of pupil dilation from each EEG power band (**Fig. 4c**). The classifier using delta and theta power performed the worst, with average accuracies of approximately 70% and 65%, respectively, indicating moderate similarity between features in their power traces during the two types of pupil dilation. All other power bands followed an increasing accuracy gradient with alpha and beta being around 80%, low gamma at 82%, and high gamma around 87% (i.e., the most differentiated power band across both dilations). This approach confirmed distinct features in the dynamics of EEG power associated with spontaneous pupil-linked arousal and pupil-linked arousal evoked by LC activation.

### Noradrenergic manipulation alters spontaneous pupil and EEG dynamics

In order to further assess the role of the LC-NE system in mediating the coupling between pupil-linked arousal and cortical state, we evaluated the effects of three FDA-approved adrenergic medications, i.e., clonidine (α-2 agonist), propranolol (β antagonist), and phentolamine (α antagonist), on EEG signals, pupil dilations, and pupil-cortical state coupling. To understand how each drug modulated cortical EEG, we first visualized the EEG power spectral changes across pharmacological manipulation and control sessions (**Fig. 5a**). All three pharmacological manipulations increased EEG power across all bands, most notably in Clonidine, implying increased oscillatory activity (**Supplemental Fig. 3**). When assessing the distribution of EEG power over the frequency bands, we noticed propranolol induced significant decreases in theta (p=0.0085, one-way ANOVA post-hoc Tukey, **Fig. 5b**), alpha (p=1.33e-06, one-way ANOVA post-hoc Tukey, **Fig. 5b**), and beta (p=0.018, one-way ANOVA post-hoc Tukey, **Fig. 5b**) power. phentolamine decreased alpha (p=0.0009, one-way ANOVA post-hoc Tukey, **Fig. 5b**) power, while clonidine resulted in very significant decreases in low gamma (p=0.005, one-way ANOVA post-hoc Tukey, **Fig. 5b**) and high gamma power (p=0.0019, one-way ANOVA post-hoc Tukey, **Fig. 5b**). Furthermore, we looked to characterize the frequency of spontaneous phasic pupil dilations across sessions for each adrenergic receptor manipulation group (**Fig. 5c**). Interestingly, when compared to the control session’s mean inter-dilation interval (43.08 s), both propranolol (85.39 s) and clonidine (78.87 s) significantly slowed the occurrence of spontaneous phasic pupil dilations, while phentolamine (66.01 s) only moderately reduced it. We next compared spontaneous phasic pupil dilations between control and manipulation sessions (**Fig. 5d**). Clonidine group notably had a significantly larger baseline pupil size (p=0.0082, one-way ANOVA post-hoc Tukey, **Fig. 5e**) and significantly reduced pupil dilation (p=1.53e-04, one-way ANOVA post-hoc Tukey, **Fig. 5e**) compared to control sessions, consistent to previous studies reporting mydriasis in rodents (Gherezghiher and Koss, 1979; Heal et al., 1989). Both phentolamine and propranolol manipulations slightly elevated baseline pupil size and reduced pupil dilations compared to control conditions, however, these differences were not statistically significant,

**Figure 5:**
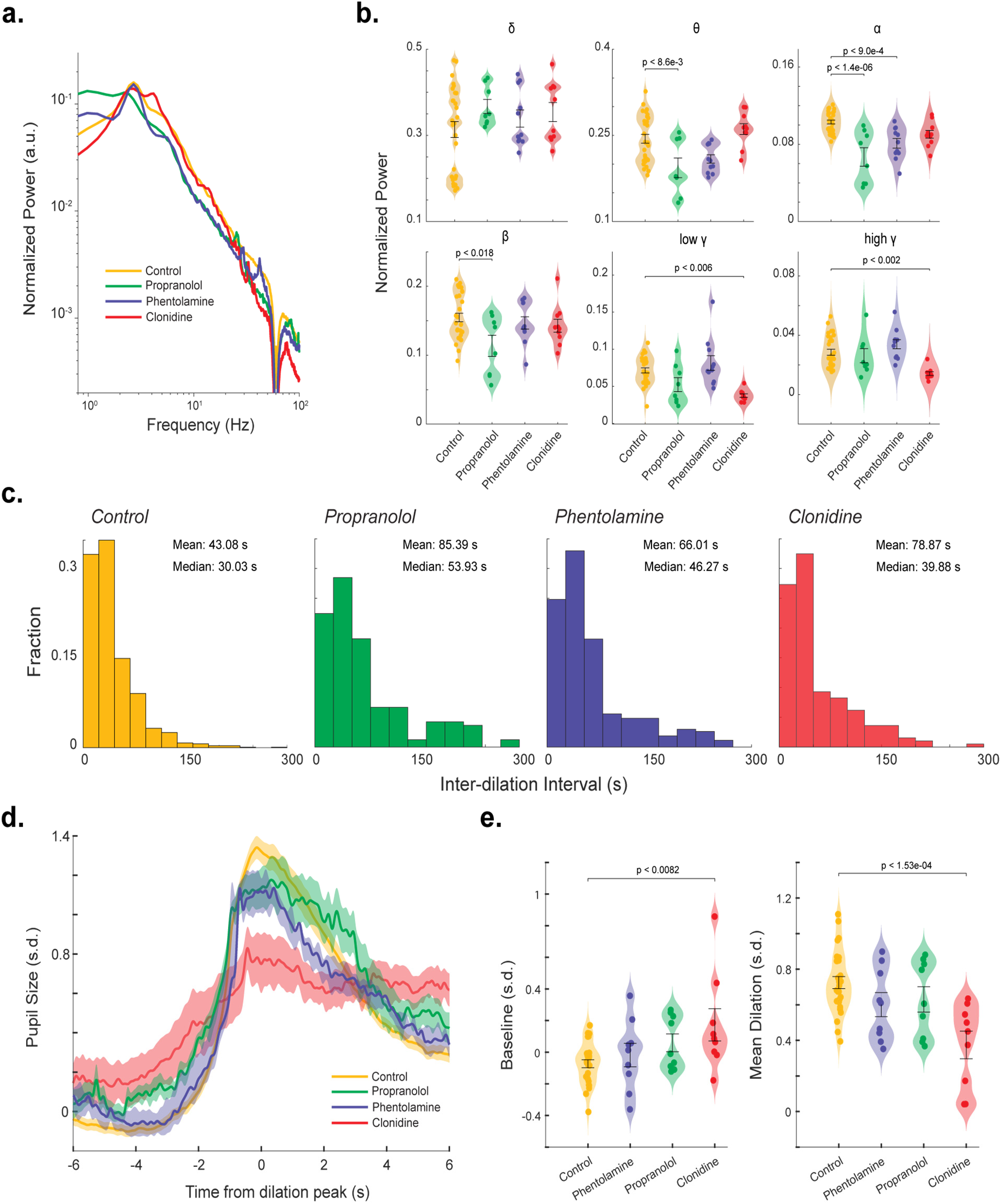
Noradrenergic manipulation alters spontaneous pupil dilations and EEG spectrum. **a.** Normalized EEG power spectral density across all noradrenergic manipulation sessions. **b.** Normalized EEG spectral power across each frequency band for all manipulation groups. **c.** Histograms of inter-dilation intervals for spontaneous phasic pupil dilations. **d.** Spontaneous pupil dilation across each manipulation and control group, aligned to peak pupil dilation. **e.** Baseline pupil size and mean pupil dilation for each manipulation and control group.

### Adrenergic manipulation affects pupil-cortical state coupling during both spontaneous and LC stimulation-evoked phasic pupil dilation

After comparing spontaneous phasic pupil dilations under different manipulations of adrenergic receptors, we examined the extent to which these pharmacological manipulations affect pupil dilations evoked by LC stimulation. LC stimulation induced smaller pupil dilations in the presence of all pharmacological manipulations compared to control sessions (**Fig. 6a**). Moreover, we observed significant disruptions in low frequency EEG power dynamics by the manipulations. During sessions with the pharmacological manipulations, we found that dynamics of EEG power in delta, theta, alpha, and beta bands were barely separatable during spontaneous and LC stimulation-evoked pupil dilations, when compared to control sessions. In addition, power in the low and high gamma bands appeared to be amplified by phentolamine during spontaneous pupil dilation, whereas power in both bands decreased during LC stimulation-evoked pupil dilation compared to control sessions.

**Figure 6:**
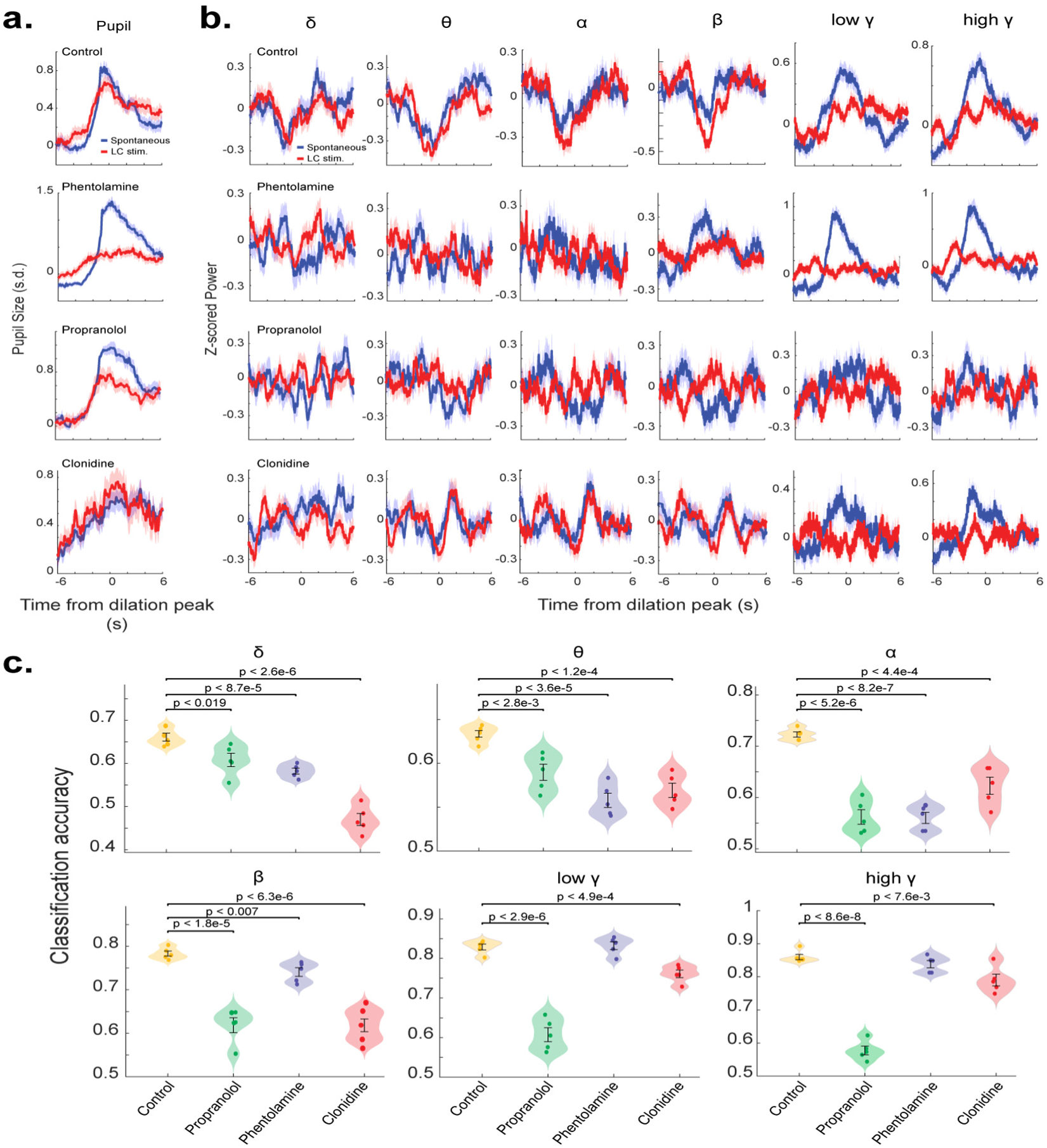
Noradrenergic manipulation reduced the accuracy of the CNN classifier in distinguishing LC stimulation-evoked from spontaneous pupil dilations based on EEG signals. **a.** Spontaneous and LC stimulation-evoked pupil dilations for the control group and each noradrenergic manipulation group. **b.** EEG power dynamics during spontaneous and LC stimulation-evoked pupil dilations for the control group and each manipulation group. **c.** Accuracy of the CNN classifier in distinguishing LC stimulation-evoked from spontaneous pupil dilations based on power dynamics of each EEG frequency band.

To quantify differences in EEG features associated with spontaneous and LC stimulation-evoked pupil dilation under these pharmacological manipulations, we again employed a CNN classifier to classify the type of pupil dilations from EEG power dynamics (**Fig. 6b**). As expected, the CNN classifier using low and high gamma power achieved comparable accuracy under phentolamine manipulation and control conditions. However, the classifier using alpha, beta, low gamma, or high gamma power exhibited the lowest accuracy during propranolol manipulation, compared to both phentolamine and clonidine conditions. Taken together, these results indicate that all three subtypes of adrenergic receptors play a critical role in modulating the coupling between pupil-linked arousal and cortical state.

## Discussion

In the present study, we investigated the extent to which the LC-NE system contributes to pupil-linked arousal coupling to cortical state. This question is grounded in the critical role of the LC-NE system in regulating brain-wide arousal levels and influencing neural information processing (Berridge and Waterhouse, 2003; Eschenko et al., 2012; Rodenkirch et al., 2019; Rodenkirch et al., 2022; Glennon et al., 2023; Ghosh and Maunsell, 2024) . Pupil diameter has become a valuable, non-invasive readout of internal arousal state and, more specifically, is tightly correlated with LC-NE activity (Joshi et al., 2016; Reimer et al., 2016; Liu et al., 2024) . While prior literature has provided functional evidence that direct activation of the LC can evoke both pupil dilations and concordant EEG power changes (typically reflecting cortical desynchronization) (Berridge and Foote, 1991; Carter et al., 2010), similar fluctuations in pupil size and cortical state also occur spontaneously (Yoss et al., 1970; Reimer et al., 2014; McGinley et al., 2015; Reimer et al., 2016; Pfeffer et al., 2022). However, a direct comparison quantifying how the relationship between cortical state during direct LC-evoked versus spontaneous arousal fluctuations has remained unexplored. Our study addresses this gap and provides new evidence supporting a distinct difference in cortical state during spontaneous phasic arousal and LC stimulation-evoked phasic arousal. Moreover, our pharmacological manipulations of adrenergic receptors reveal the importance of each subtype of adrenergic receptors in mediating cortical oscillations and their coupling to pupil size, suggesting nuances in how NE release from phasic LC activation impacts cortical dynamics compared to endogenous arousal shifts.

A critical aspect of our study is the comparison between pupil dilations evoked by direct LC phasic stimulation versus those arising spontaneously, as these events likely engage distinct underlying neural circuits and neuromodulatory contexts, potentially leading to differential coupling with cortical activity. Pupil dilations evoked by LC activation are primarily driven by a rapid and phasic release of NE that influences autonomic control pathways, including projections involving the Edinger-Westphal nucleus (EWN) complex (Breen et al., 1983; Samuels and Szabadi, 2008; Liu et al., 2017). Optogenetic LC stimulation bypasses the complex integration of afferent input from broader brainstem nuclei and forebrain regions that normally shape LC firing, resulting in pupil responses that are precise and temporally locked to LC activation. In contrast, spontaneous phasic pupil-linked arousal may arise from a distributed neuromodulatory network, likely with inputs from other regions such as the prefrontal cortex, hypothalamus, and superior colliculus (Wang and Munoz, 2015; Schneider et al., 2016). This network collectively regulates the activity of the autonomic and neuromodulatory systems, including the LC itself, often resulting in larger pupil dilations and reflecting complex, ongoing cognitive and arousal states. Despite these mechanistic differences, our results indicate that changes in cortical oscillations align temporally with pupil dilation during both types of heightened arousal, consistent with findings in previous studies in non-human primates and rodents (Ebitz and Platt, 2015; Joshi et al., 2016; Reimer et al., 2016). Specifically, we observed that during phasic arousal either occurred spontaneously or evoked by LC stimulation, EEG oscillatory power shifted from low-frequency bands to higher-frequency bands – a transition indicative of a rapid cortical sate shift associated with heightened arousal (Carter et al., 2010; Sara and Bouret, 2012). Notably these power shifts consistently preceded pupil dilations, confirming the temporal lag between cortical state fluctuations and the peripheral autonomic response (Nieuwenhuis et al., 2011; Reimer et al., 2014; Pfeffer et al., 2022). However, the context for the EEG power shifts differs: spontaneous phasic pupil-linked arousal may involve the contribution of additional neuromodulatory inputs. Serotonergic neurons from the dorsal raphe nucleus, whose phasic activity has been shown to influence pupil size (Cazettes et al., 2021; Maheu et al., 2025), modulate arousal and autonomic tone, potentially acting independently or synergistically with the LC-NE system (Pickel et al., 1977; Szabo and Blier, 2001; Maheu et al., 2025). Similarly, cholinergic input via acetylcholine from the basal forebrain is crucial for regulating cortical excitability and driving transitions between synchronized and desynchronized cortical states (Goard and Dan, 2009; Pinto et al., 2013; Lin et al., 2015; Xu et al., 2015). Notably, the cortical cholinergic activity has been shown to correlated with fluctuation of pupil size (Nelson and Mooney, 2016; Reimer et al., 2016; Liu et al., 2024). Therefore, while the sequence of cortical activation preceding pupil dilation appears generalizable, the distinct underlying circuitry and broader neuromodulatory context (including NE, 5-HT, ACh, and potentially others) associated with spontaneous vs LC stimulation-evoked phasic arousal likely contribute to the specific differences in pupil-cortical state coupling revealed in our study. Future work investigating these differences could elucidate the precise contribution of each neuromodulatory signaling to cortical arousal state relative to more globally integrated arousal processes.

Norepinephrine exerts complex modulatory control over brain functions via distinct adrenergic receptor subtypes (α-1, α-2, and β), each coupled to different intracellular signaling pathways and exhibiting unique distributions (Ramos and Arnsten, 2007; O’Donnell et al., 2012; Slater et al., 2022). To dissect the contribution of α-adrenergic receptors to NE’s influence on cortical state, we employed phentolamine, an α antagonist (Rodenkirch et al., 2019). This administration seemingly disrupted EEG oscillations in the delta, theta, alpha, and beta bands during both spontaneous and LC stimulation-evoked pupil dilation periods. However, power in both low and high gamma bands was enhanced during spontaneous pupil dilations, whereas it was disrupted during LC stimulation-evoked pupil dilations. While consistent with previous studies showing that α adrenergic receptors mediate NE’s influence on cortical state (Berridge and Waterhouse, 2003; Berridge and Spencer, 2016; Ma et al., 2024), our findings provide new evidence that α adrenergic receptor signaling constitutes a necessary pathway specifically linking phasic NE release to the modulation of cortical circuits underpinning slow oscillations. The clear distinction between gamma power during LC stimulation-evoked pupil dilations and spontaneous pupil dilations suggests that neurotransmitters other than NE may contribute more significantly to gamma oscillations (**Figure 4a**).

Interestingly, blocking α adrenergic receptors with phentolamine appears to amplify EEG gamma power during spontaneous phasic arousal. This could be due to the interplay between adrenergic receptors and other neurotransmitter receptors. For example, previous work has shown that the activation of α-2 adrenergic receptors could inhibit acetylcholine release by reducing calcium influx at cholinergic terminals (Boehm and Huck, 1995). Supporting this notion, administration of clonidine, an α-2-adrenergic agonist, resulted in a reduction of gamma power during spontaneous phasic arousal. α-2 adrenergic receptors modulate neural circuit dynamics primarily through their inhibitory nature as Gi protein coupled receptors (GPCRs). They are densely expressed in LC neurons and their terminals (Marwaha and Aghajanian, 1982; De Sarro et al., 1987; Chamba et al., 1991). Activation of these Gi GPCRs hyperpolarizes LC neurons and suppresses NE release, thereby exerting powerful control over central noradrenergic tone and consequently, arousal and cortical network activity (Hein et al., 1999; Berridge and Waterhouse, 2003). Moreover, the expression of α-2 adrenergic receptors in the EWN, which mediates the control of pupil size by the LC in the parasympathetic pathway (Liu et al., 2017). Consistent with Clonidine’s reported effects, we observed changes in pupil dynamics linked to spontaneous phasic arousal and LC stimulation, including larger baseline pupil size and longer inter-dilation intervals (Gherezghiher and Koss, 1979; Koss and Christensen, 1979; Hou et al., 2005). The inhibition of the EWN by clonidine slowed down phasic pupil dilation evoked by LC stimulation (**Figure 6a**). In addition, both amplitude and speed of spontaneous phasic pupil dilation were significantly diminished during clonidine conditions. This suggests that the other neuromodulatory nuclei mediating spontaneous phasic pupil dilations control pupil size primarily through the parasympathetic EWN. Most notably, clonidine resulted in a disruption in the coupling between spontaneous pupil-linked arousal and low-frequency EEG activity. This aligns with literature showing a reduced noradrenergic tone via α-2 agonism, leading to increased cortical synchrony, and therefore low-frequency power (Florio et al., 1975; Pastel and Fernstrom, 1984; Riekkinen Jr et al., 1990). Our results extend this finding by demonstrating that the dampening of the NE system effectively eliminates the relationship between phasic NE release and low-frequency cortical EEG activity. This decoupling could take effect by suppressing presynaptic NE to levels insufficient to reliably drive or coordinate downstream low frequency cortical network activity.

Another interesting finding is that the manipulations of α adrenergic receptors mostly disrupted EEG power during spontaneous phasic pupil dilations in the delta, theta, alpha, and beta bands, but not in gamma bands (**Figure 6a**). This suggests that the central pupil-linked arousal system modulates gamma oscillations not through alpha adrenergic receptors. However, when blocking β adrenergic receptors, EEG power across all bands was disrupted. As Gs protein coupled receptors, β adrenergic receptors activate adenylyl cyclase and downstream cAMP/PKA signaling pathways (Molinoff, 1984; Johnson, 1998). This intracellular cascade is well-known to enhance neuronal responsiveness, facilitate synaptic plasticity such as long-term potentiation (Gelinas and Nguyen, 2005; Zhang et al., 2013; Hagena et al., 2016; Borodovitsyna et al., 2017), and contribute to sustained arousal, inhibitory response, and attentional processes (Pattij et al., 2012; André et al., 2015). Furthermore, β adrenergic receptors are implicated in neural desynchronization and affect high-frequency oscillations, which are thought to reflect improved cortical processing and information transmission (Roubicek, 1976; Hazra et al., 2012). The broad disruption of EEG power by propranolol administration, particularly in high frequencies, aligns with the known effect of propranolol on high-frequency network activity (Orzack et al., 1973; Haggerty et al., 2013). The impact β adrenergic receptor inhibition across all frequency bands, contrasting with α adrenergic receptor inhibition, might reflect the widespread distribution of β adrenergic receptors throughout brain structures (Wanaka et al., 1989; Cox et al., 2008), or the engagement of downstream signaling cascades which can broadly amplify NE’s modulatory influence on overall network excitability (Waterhouse et al., 1982). Taken together, our results highlighted the critical role of adrenergic receptors in maintaining the NE-driven coordination of cortical networks that underlies robust pupil-cortical coupling during arousal events. Future work is warranted to test the role of other neurotransmitter receptors, including nicotinic and muscarinic cholinergic receptors, dopaminergic receptors, and serotoninergic receptors, in mediating the coupling between pupil-linked arousal and cortical state.

## Acknowledgements

This work was supported by AFOSR FA9550-22–1-0337, NIH R01NS119813, NIH R21MH125107 and NIH R01 MH112267. Any opinions, findings, and conclusions or recommendations expressed in this material are those of the authors and do not necessarily reflect the views of the United States Air Force.

## Competing interests

Q.W. is the co-founder of Sharper Sense, a company developing methods of enhancing sensory processing with neural interfaces. No other authors have any conflicts of interest, financial or otherwise, to disclose.

## Author Contributions

Q.W. conceived and designed research; E.W., Y.L. performed experiments and analyzed data; E.W, Q.W. interpreted results and wrote the manuscript; Q.W., E.W, Y.L. all approved the final version of manuscript. Q.W. supervised the study.

## Supplemental Figures

**Supplemental Figure 1.**
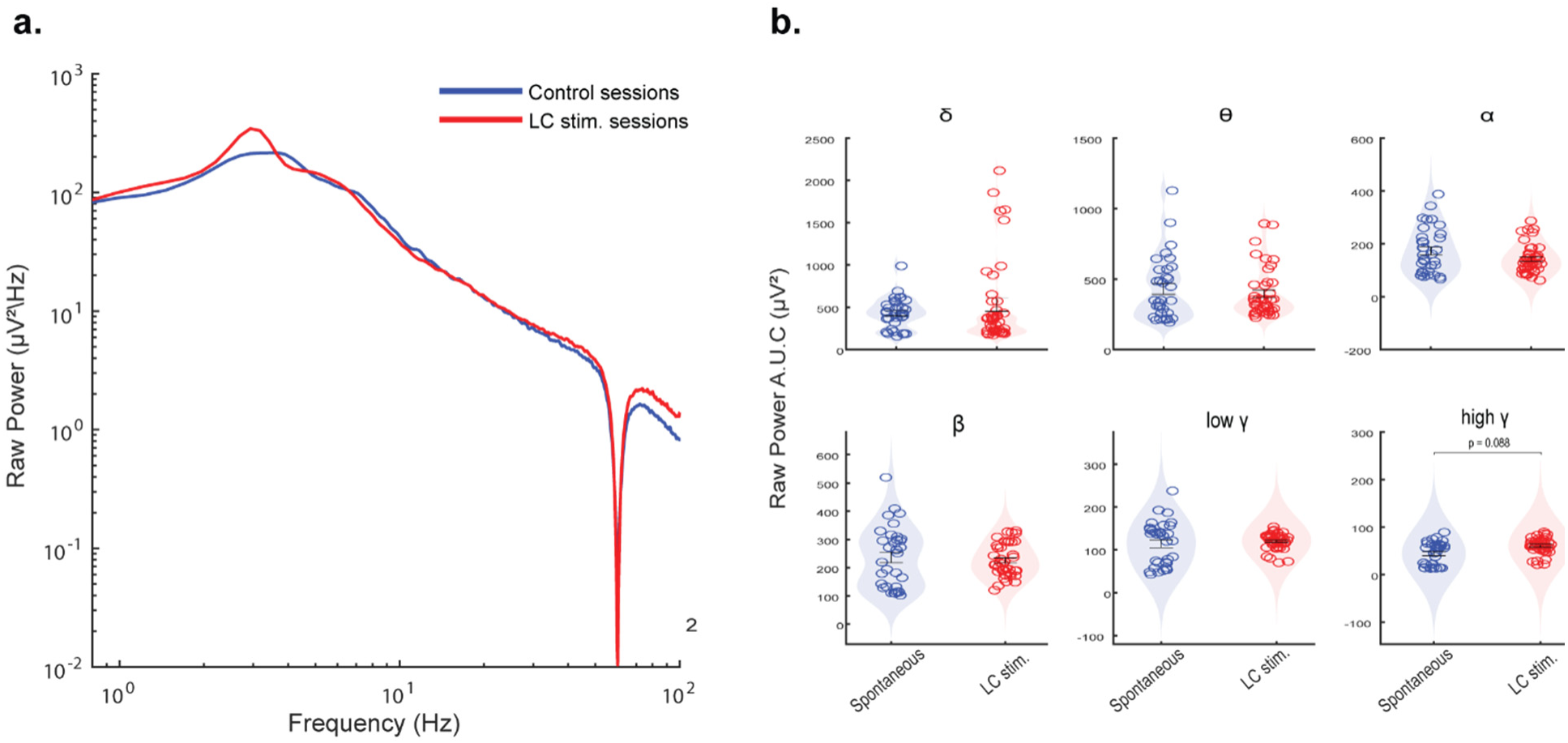
Raw EEG spectral power for spontaneous and stimulation sessions. **a.** Raw power spectral density between LC stimulation and spontaneous sessions **b.** Raw spectral power across each frequency band represented as the A.U.C.

**Supplemental Figure 2.**
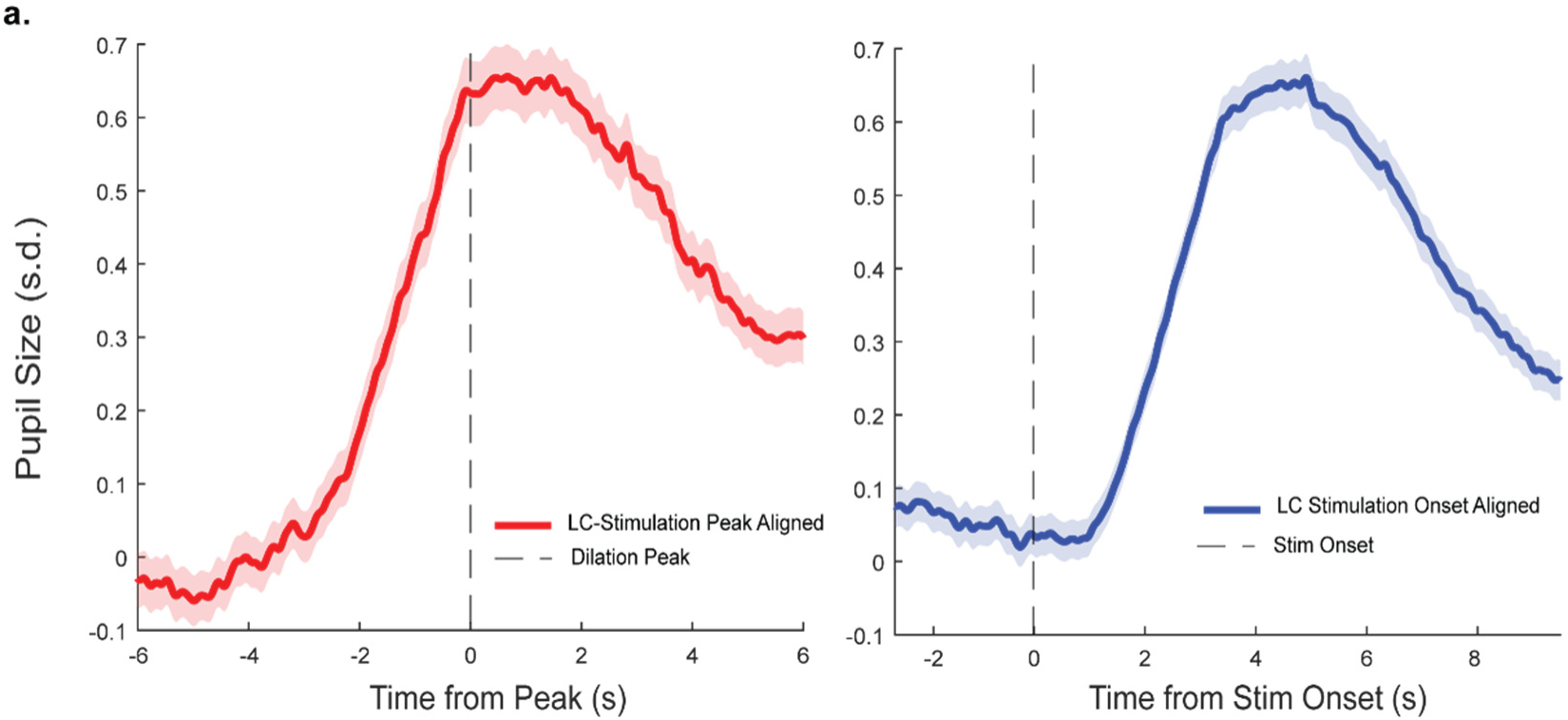
**Comparison of LC stimulation dilations aligned to peak versus alignment to stimulation onset**

**Supplemental Figure 3.**
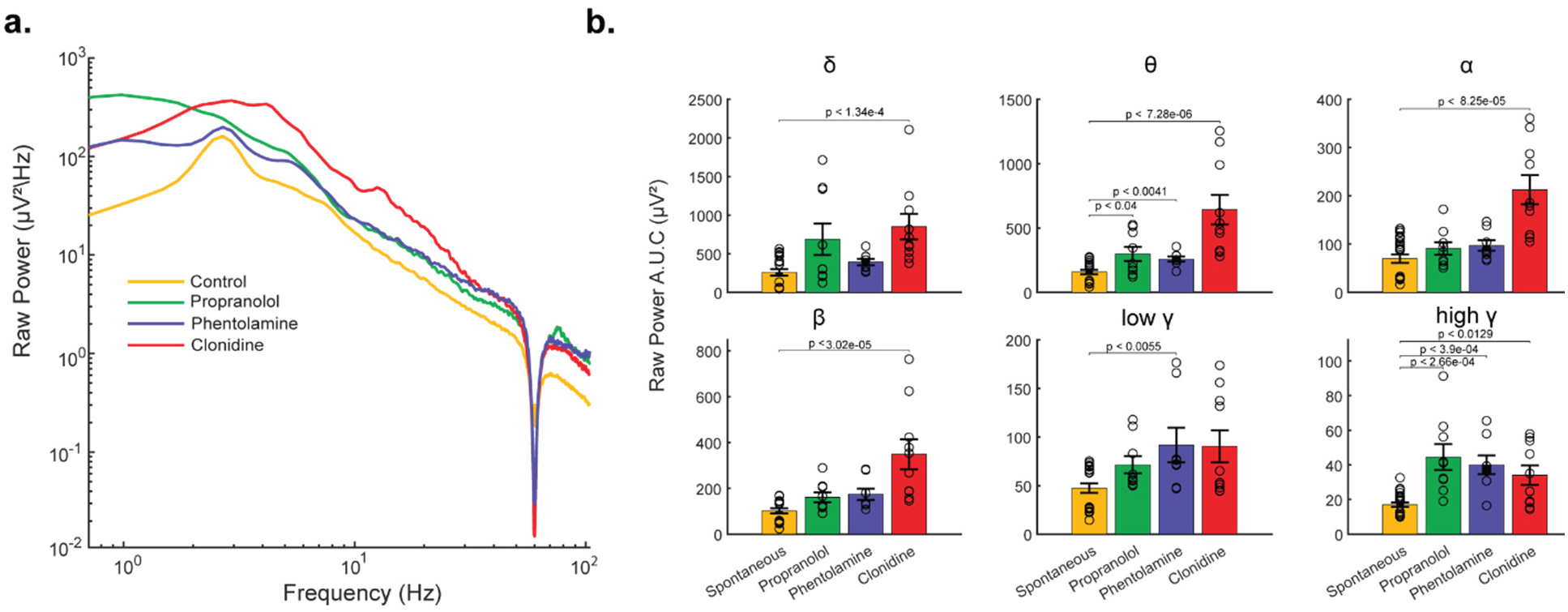
Raw EEG spectral power during noradrenergic manipulations. **a.** Raw power spectral density in control, propranolol, phentolamine, and clonidine manipulations. **b.** EEG power across each frequency band in control, propranolol, phentolamine, and clonidine manipulations.

